# A Growth-Based Framework for Scaling Infant Anthropometry in Motion Capture Studies

**DOI:** 10.64898/2026.05.27.728208

**Authors:** Ketaki Inamdar, Nishat Tasnim Koli, Sara Gamal, Valeriya Gritsenko

## Abstract

Biomechanical analysis of infant motor behavior is complicated by rapid growth and large inter-subject variability in body size and segment proportions. We present a growth-adjusted anthropometric scaling framework that integrates Centers for Disease Control and Prevention (CDC) growth charts with three-dimensional motion capture to estimate subject-specific infant biomechanics during prone play. Nineteen infants aged 2–6 months were recorded during spontaneous prone movement. Weight-for-age percentiles were computed using CDC LMS parameters and used to infer body length from length-for-age references, ensuring internal consistency between weight and length estimates. Body segments were scaled using published infant proportions and validated against head and shoulder widths measured from motion capture. Infant head and trunk proportions were substantially larger than adult values, whereas limb proportions were similar. Estimated and measured dimensions of head and trunk width showed good agreement, with mean root mean squared errors of approximately 3 cm. Adult-based scaling produced head and trunk length errors of up to 6 cm. This growth-adjusted framework improves anatomical fidelity and enables normalization of biomechanical measures across infants with differing growth trajectories.

## I. Introduction

Early infancy is characterized by rapid changes in body mass, length, and segment proportions, occurring concurrently with the emergence of foundational motor skills such as head control, trunk stabilization, and upper-limb support. Prone play, commonly referred to as tummy time, provides a critical context for observing and quantifying these early motor behaviors [1].

Quantitative analysis of motor behavior during early infancy presents unique challenges due to rapid and highly variable growth. In contrast to adult populations, for which extensive anthropometric databases exist [2], infant body dimensions change substantially over short time scales, and considerable variability persists even among infants of the same chronological age. Consequently, biomechanical analyses that rely solely on age-based grouping risk conflating differences in neuromuscular control with differences in growth status and body proportions.

Although standardized growth charts from the Centers for Disease Control and Prevention (CDC) and World Health Organization (WHO) are widely used in clinical settings to characterize infant growth, these tools are not routinely integrated into biomechanical data processing pipelines. The objective of this study is to develop a growth-adjusted scaling framework that explicitly integrates CDC growth chart data into infant biomechanical analysis and to estimate subject-specific growth percentiles and body length using the LMS statistical method. In this method, L denotes the Box–Cox power accounting for distribution skewness, M denotes the median value at a given age, and S denotes the coefficient of variation.

## II. Methods

### A. Participants and Experimental Protocol

Nineteen infants (age range: 2–6 months) participated in the study; demographic characteristics are summarized in Table I. Six participants were born preterm, with gestational ages ranging from 23 to 36 weeks. All experimental procedures were approved by the Institutional Review Board of West Virginia University (protocol #2306804216), and written informed consent was obtained from a parent or legal guardian prior to participation.

**TABLE I.** Participant Demographics and anthopometric measurements.

| ID | Sex | Age (adjusted age, mo) | Measured weight (kg) | Inferred height (m) | Shoulder width (m) | Head width (m) |
| --- | --- | --- | --- | --- | --- | --- |
| C1 | m | 6 | 9.98 | 0.723 | 0.179 | 0.149 |
| C2 | f | 4 | 6.8 | 0.63 | -<sup>a</sup> | 0.15 |
| C3 | m | 4 | 6.8 | 0.632 | 0.161 | 0.146 |
| C4 | m | 4 | 6.28 | 0.617 | 0.141 | 0.143 |
| C5 | f | 3 | 5.1 | 0.577 | 0.169 | 0.148 |
| C6 | m | 6 | 7.7 | 0.667 | 0.176 | 0.161 |
| C7 | m | 5 | 7.6 | 0.659 | 0.161 | 0.152 |
| C8 | f | 6 | 7.26 | 0.653 | 0.182 | 0.144 |
| C9 | m | 3 | 5.5 | 0.588 | 0.148 | 0.12 |
| C10 | m | 7 | 6.68 | 0.649 | 0.174 | 0.148 |
| C11 | m | 3 | 5.55 | 0.592 | 0.152 | 0.121 |
| C12 | m | 4 | 4.35 | 0.554 | 0.176 | 0.103 |
| C13 | m | 6 | 4.68 | 0.578 | 0.189 | 0.152 |
| P1 | f | 5 (4) | 5.08 | 0.578 | 0.121 | 0.144 |
| P2 | m | 7 (3) | 6.12 | 0.637 | 0.149 | 0.149 |
| P3 | f | 8 (7) | 9.53 | 0.713 | 0.187 | 0.145 |
| P4 | f | 5 (5) | 4.99 | 0.581 | 0.193 | 0.141 |
| P5 | f | 7 (5) | 4.99 | 0.581 | 0.173 | 0.139 |
| P6 | f | 10 (7) | 5.52 | 0.607 | 0.137 | 0.133 |

<sup>a</sup>. Incorrect marker placement.

Infants were positioned prone on a padded surface and allowed to engage in spontaneous movements during free play. Motion was recorded using an active-marker motion capture system (Impulse, PhaseSpace Inc., San Leandro, CA, USA). Markers were placed on predefined anatomical landmarks of the head and trunk (Fig. 1). Marker trajectories were sampled at 480 Hz. Recording sessions lasted between 1.5 and 8.8 minutes. Data acquisition was paused whenever an infant exhibited signs of discomfort, at which point the caregiver provided soothing. Recording resumed once the infant was calm; if distress persisted, the session was terminated. Additionally, participant’s weight, sex assigned at birth, and gestational age was recorded.

**Fig. 1.**
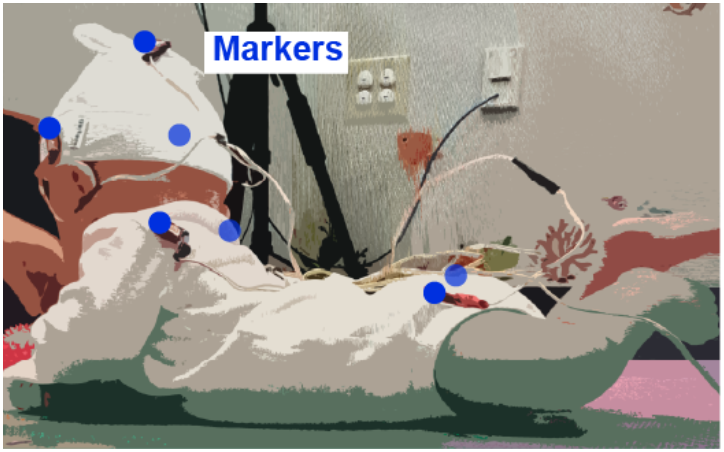
Position of participant and marker placement.

The motion analysis relies on inverse kinematics. However, accurate joint angle calculations rely on anthropometry (segment lengths of thigh, trunk, etc.) usually inferred from height. Since it is difficult to estimate infant true height, we used data from the CDC to infer height from weight. We used the CDC-provided growth chart data files for weight-for-age and length-for-age. These data sets include age-specific LMS parameters, where L represents the Box–Cox power accounting for distribution skewness, M represents the median value at a given age, and S represents the coefficient of variation [3], [4].

The LMS formulation is useful for continuous conversion between raw anthropometric measurements, z-scores, and percentiles. Age in the CDC growth charts is reported at half-month intervals representing the midpoint of each month. For each subject, recorded age was mapped to the appropriate half-month age prior to interpolation of LMS parameters.

### B. Anthropometric Estimation and Inverse Kinematics Scaling

Inverse kinematics requires subject-specific anthropometric parameters to compute joint angles accurately [5]. While infant body length is commonly obtained in clinical settings, direct measurements can be affected by posture, soft tissue compression, and examiner-dependent variability, which may introduce error when used for biomechanical scaling. Therefore, body length was estimated indirectly from body weight using normative growth data. Specifically, CDC growth chart data files for weight-for-age and length-for-age were used, which provide age- and sex-specific LMS parameters describing the distribution of anthropometric measures [3], [4].

Subject-specific weight percentiles were computed from measured body weight, age in months, and sex. LMS parameters (L, M, S) were linearly interpolated from the CDC weight-for-age tables at the mapped age. The measured weight (X) was converted to a z-score (Z) using the standard LMS transformation:

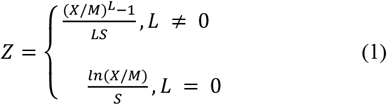

where X denotes the observed measurement and L, M, and S are the interpolated LMS parameters. The corresponding percentile was obtained by evaluating the cumulative distribution function of the standard normal distribution at Z. This procedure yields percentile estimates consistent with CDC growth standards.

Estimated body length was obtained by applying the subject’s weight percentile to the CDC length-for-age distribution for the same age and sex. LMS parameters for length-for-age were interpolated at the mapped age. The percentile was converted to a z-score and then transformed to an estimated body length using the inverse LMS equation. This approach ensures that the estimated length corresponds to the same relative position within the growth distribution as the subject’s body weight, thereby maintaining internal consistency across anthropometric measures.

### C. Body Segment Scaling

Estimated total body length was used to scale individual body segments according to published anthropometric proportions for infants < 6 months old [2]. Segment geometries were modeled as rectangular prisms (chest, pelvis, hands, and feet), an ellipsoid (head), and cylinders (upper and lower arm and leg segments) (Fig. 2). The distal forearm segments were split using 2/3 and 1/3 fractions to accurately represent pronation/supination degrees of freedom [6]. Similarly, the distal leg segments were split to represent internal/external rotation of the foot using ½ fractions. Segment lengths were defined as fixed fractions of total body length based on reported proportional data. The trunk and pelvis girth was estimated from the published infant chest circumference measurements [7]. Cylinder diameters and hand and feet prism girth dimensions were estimated empirically.

**Fig. 2.**
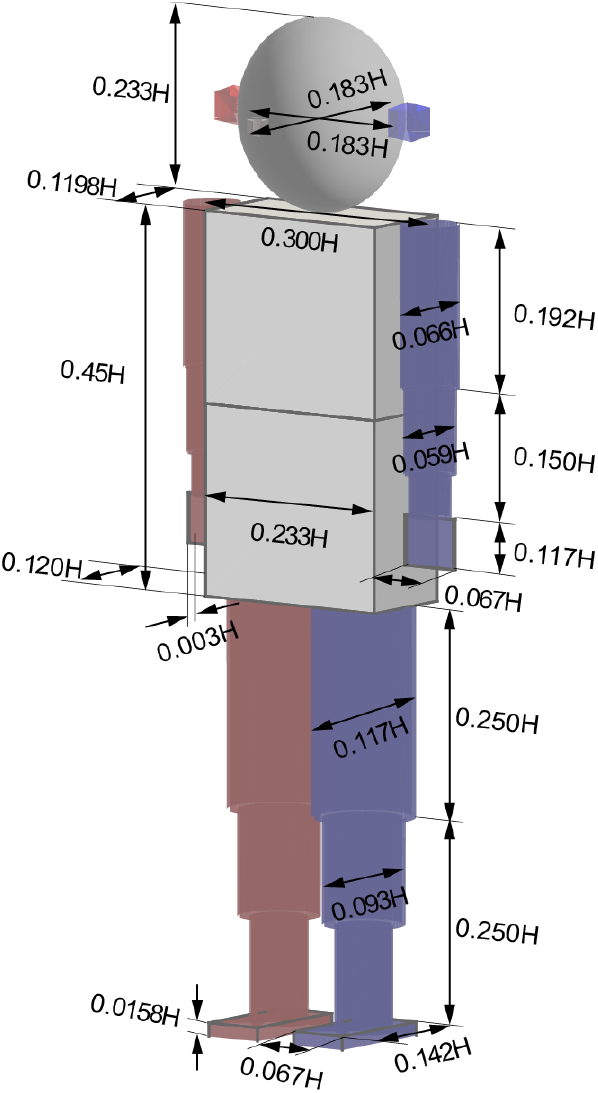
3D model illustrating segment shapes and proportions. H denotes total height top of the head to bottom of feet.

Validation of inferred segment dimensions was performed using head width and shoulder width measurements extracted from the motion capture recordings. Estimated and measured dimensions were compared using the root mean squared error (RMSE) metric.

## III. Results

The CDC–WHO growth curves for male and female infants begin to diverge at approximately 6 months of age (Fig. 3).

**Fig. 3.**
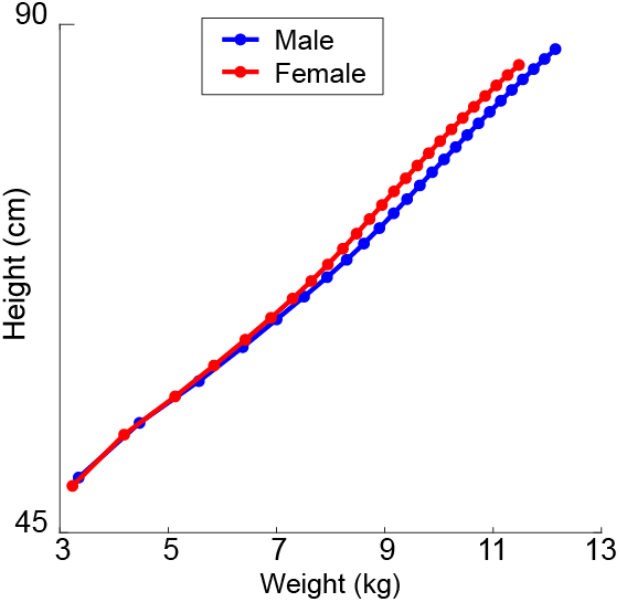
Example of CDC grouth charts for average (50%) males and females over the first 24 months of life.

Accordingly, a single set of anthropometric segment proportions was applied to all infants younger than 6 months, independent of sex. The resulting proportions are summarized in Table II.

**TABLE II.** Height-Normalized Body Segment Proportions for Infants (<6 Months, 50TH Percentile)

| Segment | Shape | Length / Height | Width / Height | Girth / Height |
| --- | --- | --- | --- | --- |
| Head | ellipsoid | 0.233 | 0.183 | 0.183 |
| Trunk | box | 0.225 | 0.3 | 0.1198 |
| Pelvis | box | 0.225 | 0.233 | 0.1198 |
| Humerus (upper arm) | cylinder | 0.192 | 0.066 | 0 |
| Radius/Ulna (forearm) | cylinder | 0.15 | 0.0594 | 0 |
| Hand | box | 0.117 | 0.067 | 0.0033 |
| Femur (thigh) | cylinder | 0.25 | 0.117 | 0 |
| Tibia (shank) | cylinder | 0.25 | 0.0932 | 0 |
| Foot | box | 0.142 | 0.067 | 0.0158229 |

Fig. 2 illustrates the three-dimensional infant body model at the 50th percentile body length (0.60 m), scaled using the inferred segment proportions. The head height proportion in infants (0.23 of total body length) was nearly twice the corresponding adult proportion (0.13) [8]. Similarly, the trunk length proportion (0.45) was substantially larger than the adult value (0.29) [8]. Consequently, scaling based on adult segment proportions produced mismatches in head and chest (upper body only) lengths ranging from 3 to 6 cm (Fig. 4A). The remaining segment proportions were comparable to adult values. These proportions reflect characteristic infant morphology, with a relatively large head and trunk and proportionally shorter limb segments. Agreement between estimated and measured segment dimensions obtained from three-dimensional motion capture was high (Fig. 4B). For infants born at 39 or 40 weeks of gestation, the mean root mean squared error (RMSE) between estimated and measured shoulder width was 2.7 cm, and the RMSE for head width was 3.2 cm. For infants born preterm, the mean RMSE values were similar, 3.1 cm for shoulder width, and 2.8 cm for head width.

**Fig. 4.**
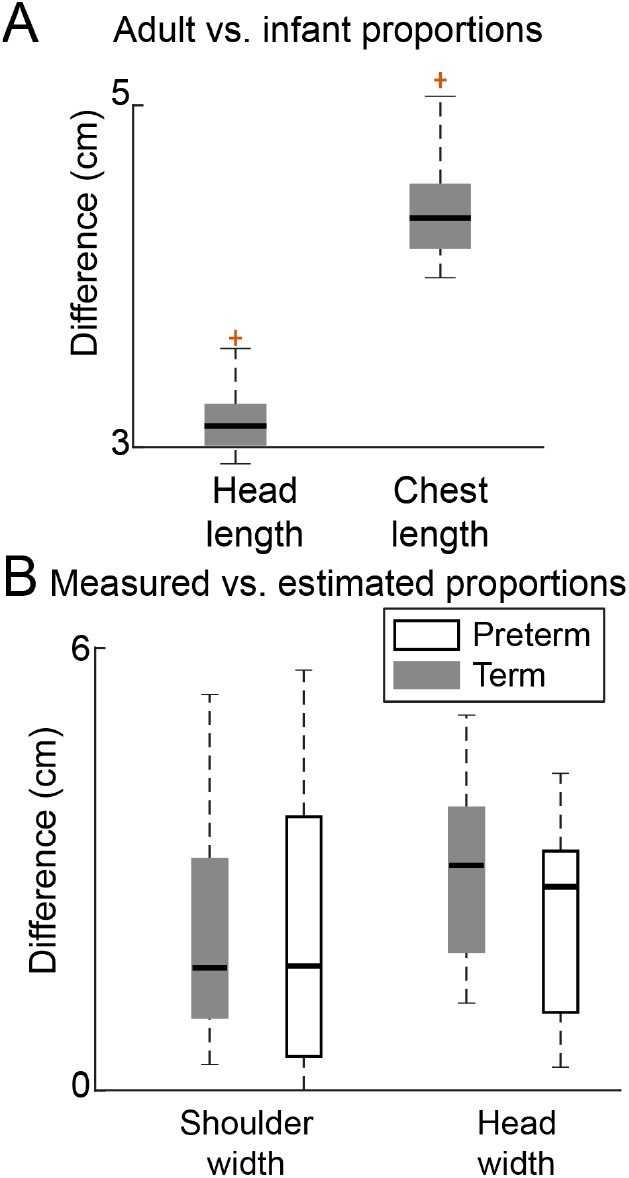
Errors in estimating segment lengths from adult proportions (A) and from CDC growth charts (B). Thick lines show median values of absolute differences, boxes show quartile ranges, dashed lines show full range of errors, red crosses show outliers.

Together, these results validate the proposed scaling approach and establish a reference anthropometric model for estimating subject-specific segment dimensions in early-infant biomechanical analyses.

## IV. Discussion

This study developed and validated a growth-adjusted anthropometric scaling framework for biomechanical analysis of infants during prone play by integrating CDC growth chart data with motion capture–based kinematics. Subject-specific weight percentiles were first estimated using CDC LMS parameters, and these percentiles were subsequently used to infer body length from length-for-age growth distributions, ensuring internal consistency between weight and length estimates. The proposed framework enables consistent, subject-specific scaling of body segments while accounting for the unique morphological characteristics of early infancy.

The CDC–WHO growth curves show that male and female growth trajectories diverge only after approximately 6 months of age. Accordingly, a single set of infant anthropometric proportions was applied to all participants younger than 6 months. The resulting segment proportions reflected characteristic infant morphology. The head height proportion was nearly twice the adult value, while the trunk length proportion exceeded the adult proportion by more than 50%. These findings align with established principles of cephalocaudal development, where growth proceeds from head to toe, with the head representing a disproportionately large fraction of total body length during infancy [9].

Validation against motion capture–derived measurements showed good agreement between estimated and measured segment dimensions. The mean RMSE values were about 3 cm for shoulder and head width across all participants. Consequently, scaling based on adult segment proportions for head and trunk produced mismatches, with errors reaching up to 6 cm. Such errors would propagate through subsequent biomechanical calculations, including joint center estimation, segment mass distribution, and moment arm calculations, potentially compromising the validity of kinematic and kinetic analyses. These results demonstrate that growth-adjusted scaling improves anatomical fidelity relative to conventional adult-based scaling and provides a feasible approach for estimating subject-specific infant anthropometry when direct length measurements are unavailable.

Several limitations should be noted. First, body length was not measured directly but inferred from weight percentiles using population-level growth references. This approach assumes that individual infants follow typical CDC growth trajectories and may be less accurate for infants with atypical growth patterns, including preterm infants or those with growth restrictions. Although, the mean errors for both groups of our participants were similar. Second, segment girths and some geometric dimensions were estimated empirically rather than measured directly, introducing potential systematic uncertainty in segment mass and inertia estimates.

Validation was limited to head width and shoulder width, which represent only a subset of segment dimensions and do not directly assess the accuracy of limb lengths, segment masses, or inertial properties critical for inverse dynamics. The sample size was modest (n = 19), and six participants were born preterm, which may introduce additional heterogeneity not explicitly modeled by the scaling procedure. Finally, the framework assumes fixed segment proportions within the 2–6 month age range and does not capture nonlinear changes in body proportions that may occur in early infancy.

Despite these limitations, the proposed growth-adjusted scaling framework provides a principled and reproducible method for incorporating standardized growth information into infant biomechanical analyses. By explicitly integrating CDC growth charts with inverse kinematics scaling, this approach enables normalization of kinematic and neuromuscular measures across infants with different growth trajectories and reduces confounding between motor control development and body size.

This framework facilitates quantitative comparisons of infant motor behavior during prone play and may improve the interpretation of developmental changes in posture, head control, and upper-limb support. More broadly, the method establishes a foundation for longitudinal studies of early motor development and for future integration of growth-adjusted biomechanical modeling into clinical assessments of neuromotor risk and developmental delay.

